# CLIPPER 2.0: Peptide level annotation and data analysis for positional proteomics

**DOI:** 10.1101/2023.11.30.569335

**Authors:** Konstantinos Kalogeropoulos, Aleksander Moldt Haack, Elizabeta Madzharova, Antea Di Lorenzo, Rawad Hanna, Erwin M. Schoof, Ulrich auf dem Keller

## Abstract

Positional proteomics methodologies have transformed protease research, and have brought mass spectrometry (MS)-based degradomics studies to the forefront of protease characterization and system-wide interrogation of protease signaling. Considerable advancements in sensitivity and throughput of liquid chromatography (LC)-MS/MS instrumentation enable generation of enormous positional proteomics datasets of natural and protein termini and neo-termini of cleaved protease substrates. However, such progress has not been observed to the same extent in data analysis and post-processing steps, which arguably constitute the largest bottleneck in positional proteomics workflows. Here, we present a computational tool, CLIPPER 2.0, that builds on prior algorithms developed for MS-based protein termini analysis, facilitating peptide level annotation and data analysis. CLIPPER 2.0 can be used with several sample preparation workflows and proteomics search algorithms, and enables fast and automated database information retrieval, statistical and network analysis, as well as visualization of terminomic datasets. We demonstrate our tool by analyzing GluC and MMP9 cleavages in HeLa lysates. CLIPPER 2.0 is available at https://github.com/UadKLab/CLIPPER-2.0.

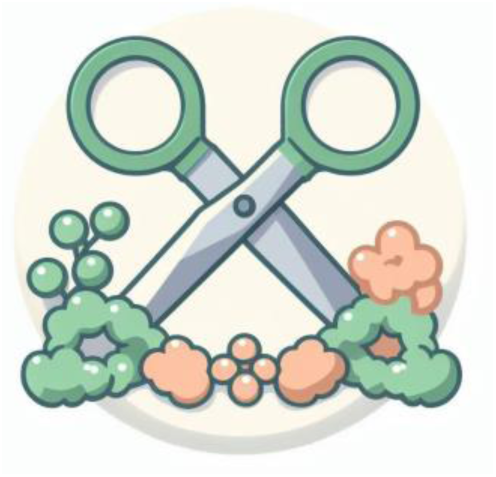

## Introduction

Mass spectrometry has been at the forefront of proteomics research for more than two decades, as it facilitates high-throughput and system-wide analysis of biomolecules^1^. Degradomics or positional proteomics, the area of proteomics pertaining to the identification of protein termini for investigation of protein processing and proteolytic activity, has undergone a massive development during this period^2,3^. With the advent of instruments capable of high resolution and throughput, as well as reliable and scalable search engines, and most importantly application-specific sample preparation workflows and enrichment strategies, it is now possible to detect thousands of natural or neo-protein N-termini from minute sample amounts^4–8^.

Despite progress in sample preparation and data acquisition, data analysis for datasets generated by degradomics workflows have not kept up with the pace of development in the rest of the analytical pipelines. Contemporary datasets are large in size, and there is a need for robust, dynamic tools for top-down overview of results, as well as detailed analysis of individual N-termini or cleavages detected. Even though there is a plethora of tools available for proteomics data analysis and visualization, adaptation to degradomics datasets is not feasible or far from optimal. This is due to the requirement for specific analyses, methods and considerations when working with proteolytic activity and peptide level datasets. Several attempts have been made to develop workflows for degradomics data analysis such as MANTI^9^ for MaxQuant^12^, Fragterminomics^11^ for Fragpipe^12^, and originally CLIPPER^13^ for the Trans Proteomic Pipeline (TPP)^14^. However, these tools are software-specific, and mostly limited to cleavage annotation of N-terminomic datasets. Therefore, there is a need for an end-to-end workflow for the analysis of datasets.

Here, we present a data analysis tool called CLIPPER 2.0, which is substantially faster than prior tools, allowing for processing of tens of thousands of peptides in minutes, and performs both annotation, statistical analysis, and creates visualizations. It accepts search results directly from the widely used software suites Spectronaut™^15^ and Proteome Discoverer™^16^, while input data can be readily adapted from other pipelines such as MaxQuant^10^ and FragPipe^12^. CLIPPER 2.0 accepts quantification data from experiments using dimethylation and TMT labeling, with data acquired either in data-dependent (DDA) or data-independent acquisition (DIA) mode^17^, with options to extend to other negative enrichment or positive enrichment strategies. CLIPPER 2.0 can be used as a standalone command line tool, or as a local browser based application that is fully compatible with Windows, and partially compatible with Mac computers.

Excluding the web app, CLIPPER 2.0 is built entirely in python, and integrates the publicly available databases to annotate observed peptides and potential protease cleavages with comprehensive information (Fig 1a). The tool offers positional annotation of identified peptides, cleavage site environments, and known cleavage event annotation, and predicts protease activity, solvent accessibility, and protein secondary structure of cleavage sites. CLIPPER 2.0 also retrieves ontology and functional information, and estimates protein localization in different cell types and tissues from available databases. In quantitative experiments, statistical tests identify peptides with significant abundance differences between conditions, and several visualizations are generated to illustrate data quality metrics and enable exploratory cleavage focused and system wide data analysis (Fig. 1b).

**Fig. 1.**
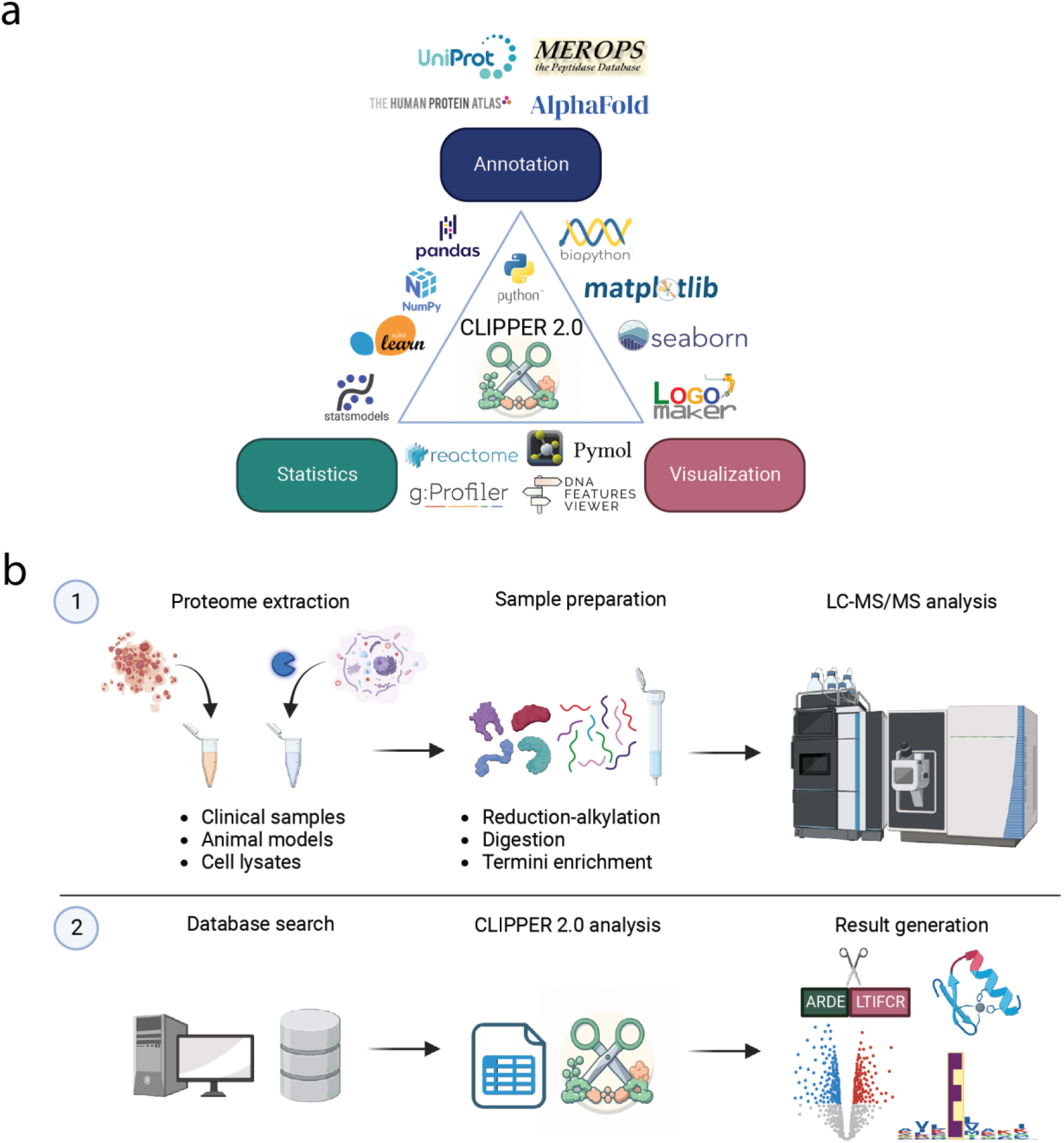
The CLIPPER 2.0 ecosystem and workflow. a) CLIPPER 2.0 integrates annotation, statistics, and visualization in a single toolkit. b) Diagram illustrating the sample preparation (1) and data flow (2).

## Methods

### Cell culture

HeLa cells were cultured with DMEM (ThermoFisher Scientific, cat. no. 11965092) until confluency, then harvested with trypsin (ThermoFisher Scientific, cat. no. 252000560) and pelleted with centrifugation at 800 x g for 5 minutes. The pellets were washed with PBS and centrifuged again. The supernatant was discarded, and the pellet was resuspended in 6 M guanidine hydrochloride (GuHCl). Cells were lysed with probe sonication (20% amplitude, 5 sec on, 5 sec off) on ice. Afterwards, the lysates were centrifuged at 4700 x g, 4 ͦ C for 15 minutes, and the supernatant was transferred to a new tube. Protein amount was quantified using Nanodrop One (ThermoFisher Scientific).

### Degradomics experiments

To demonstrate the applications of CLIPPER 2.0, we generated two novel datasets for the proteases GluC and MMP9. The GluC experiment was a TMT-TAILS experiment, while the MMP9 dataset was generated using the HUNTER workflow. Both experiments were analyzed in DDA mode.

### TAILS

In the first experiment, we started with 5 μg of proteome diluted in 20 μl of 100 mM 4-(2-hydroxyethyl)-1-piperazineethanesulfonic acid (HEPES). We used 5 biological replicates of HeLa lysates which were treated with the GluC protease (Promega, cat. no. V1651) at a ratio of 1:500 and incubated in a thermomixer for 15 minutes at 37 ͦ C. The reaction was quenched by addition of GuHCl in 2.5 M final concentration and heat denaturation. Along with 5 more control replicates, samples were reduced and alkylated with 10 mM tris(2-carboxyethyl)phosphine (TCEP) and 40 mM 2-Chloroacetamide (CAA), respectively. TMTpro reagents (ThermoFisher Scientific, cat. no. A52045) were used to label protein termini and lysines in each sample at 1:4 protein: label (w/w) ratio, incubated for 40 minutes at room temperature. The reaction was quenched with 100mM ammonium bicarbonate. Afterwards, samples were combined and SP3 cleanup^18^ was employed to remove excess reactants. The combined proteome was digested with trypsin (Promega, cat. no. V5280) at 1:50 protease: protein (w/w) at 37 ͦ C overnight. A small volume (10%) was removed as the preTAILS sample, acidified with 1% trifluoroacetic acid (TFA) and stored until further processing. To deplete tryptic peptides, the remaining sample was incubated overnight with 50mM sodium cyanoborohydride and HPG-ALD (https://ubc.flintbox.com/technologies/888fc51c-36c0-40dc-a5c9-0f176ba68293) polymer at 1:4 peptide: polymer (w/w). The resulting sample was spun down with an Amicon 30kDa filter (Merck, UFC503096) to retrieve unbound N-termini. The TAILS sample was acidified, and appropriate volumes for 500 ng injections were loaded on EvoTips with the standard loading protocol.

### HUNTER

A similar procedure was performed for sample preparation of the MMP9 experiment. The notable differences were starting with 20 μg of proteome digested with activated MMP9, the use of 60 mM formaldehyde and 50 mM sodium cyanoborohydride instead of TMTpro to label protein termini and lysines, and the depletion of tryptic peptides with undecanal instead of the HPG-ALD polymer. After SP3 cleanup, digested peptides were incubated with undecanal (Merck, cat. no. U2202) at 1:50 peptide: undecanal (w/w) at 50 ͦ C for 90 minutes. Pure EtOH and 25% TFA were added for a final concentration of 40% EtOH and 1% TFA. The solution was passed through a conditioned SepPak (Waters, cat. no. 186000308) column, and the flow through containing the enriched N-termini was collected. Both standard proteome and enriched fraction were desalted with a SoLaμ SPE plate (ThermoFisher Scientific, cat. no. 60209-001), and analyzed separately with LC-MS.

### Data acquisition and analysis

For the TMT-TAILS GluC experiment, we used the EvoSep One liquid chromatography system (Evosep, Denmark) in line with an Orbitrap Exploris 480 mass spectrometer equipped with a FAIMSpro interface. Peptides were separated with a PepSep column (Bruker Daltonics, cat. no. 1895812) over 118 minutes with the Whisper100 10SPD method, and injected to the mass spectrometer with a PepSep emitter (Bruker Daltonics, cat. no. 1893519) with a positive ion spray voltage of 2300 V, ion transfer tube temperature of 240 ͦ C. The carrier gas flow was set to 3.6 L/min, and FAIMS was operated in standard resolution in 3 different compensation voltages of -40, -55, and -70 with otherwise similar acquisition settings. MS scans were acquired in the Orbitrap at 120,000 resolution and a scan range of 375-1500, maximum injection time of 50 ms, RF Lens of 60%, and normalized AGC target of 300%. Filters of charge state between 2-7, dynamic exclusion with a duration of 60 seconds with 10 ppm mass tolerance, minimum intensity of 5000, and precursor fit at 70% were included. MS/MS scans were acquired in DDA as centroids in the Orbitrap was operated at 60,000 resolution and isolation window of 0.7 m/z, first mass at 110, HCD fragmentation at 34% NCE, maximum injection time of 118 ms, and normalized AGC target at 75%.

For the dimethyl-HUNTER experiment, an EASY-nLC™ 1200 System (ThermoFisher Scientific) was used in line with an Orbitrap Q Exactive mass spectrometer. Peptides were separated using an EASY–Spray™ column and emitter (ThermoFisher Scientific, cat. no. ES904) over 70 minutes with 10% buffer B (80% acetonitrile, 0.1% formic acid). The gradient was as follows: starting with 10% buffer B, with a linear increase to 23% until minute 43, to 38% until minute 55, 60% until minute 60, 95% until minute 63, with a steady composition to wash the column until minute 70. Peptides were ionized with spray voltage of 2000 V and transfer tube temperature of 275 ͦ C. MS scans were acquired in the Orbitrap with 70,000 resolution and a scan range of 300-1750 m/z, AGC target of 3,000,000 and maximum injection time of 50 ms. For precursor selection, a top10 DDA mode was used with additional filters of intensity threshold at 5,000 and dynamic exclusion of 30 seconds. MS/MS scans were acquired with a resolution of 17,500 and isolation window of 1.6 m/z, fixed first mass at 120, and NCE of 27% in profile mode.

The raw data from both experiments were analysed with Proteome Discoverer v2.4. The data were searched against the human proteome (Uniprot reviewed 20,311 sequences, accessed 12/07/2022) with semi-tryptic specificity and two maximum missed cleavages. For the GluC dataset, methionine oxidation (+15.995) and asparagine deamidation (+0.984) were set as dynamic modifications. TMTpro (+304.207) and acetylation (+42.011) were added as dynamic modifications on the peptide and protein terminus, respectively. Cysteine carbamidomethylation (+57.021) and lysine TMTpro labeling were added as static modifications. Peptide-spectrum matching was performed with Sequest HT, and FDR control with Percolator. Proteins were quantified using the Reporter Ion Quantifier node. The same settings were also applied to the MMP9 dataset, with the exception of light dimethylation (+28.031) modification in place of TMTpro labeling, and quantification based on MS1 precursor signal with the Minora Feature Detector and the Precursor Ions Quantifier nodes. Results were exported for processing with CLIPPER 2.0.

### Description of the CLIPPER 2.0 architecture and capabilities

CLIPPER 2.0 is an open source software tool written in Python 3^19^, a high level programming language with a large developer base, with support, maintenance and addition of new features is thus more robust when compared to existing tools. All modules are called by a central script during runtime. The user has the option of using the command line tool, or a browser-based graphical user interface (GUI) for data submission. Advanced users can also install CLIPPER 2.0 as a python package, which would enable custom workflows and easier programmatic access and integration with other tools. CLIPPER 2.0 comes with the ability to utilize threading parallelization, significantly speeding up annotation and results generation in modern computer architectures. The CLIPPER 2.0 repository is equipped with installation instructions and guides on how to use the tool, and will be maintained and expanded regularly.

The main and only required input for analysis with CLIPPER 2.0 is a peptide table containing the identified sequences, their modifications and the proteins of origin. By default, CLIPPER 2.0 recognizes the output tables and data columns from the Proteome Discoverer, Spectromine and Spectronaut search engines. However, the user can easily use peptide tables generated from other software by simply renaming the data columns containing the necessary information, or by modifying the code to recognize the desired column names. CLIPPER 2.0 accepts data coming from different acquisition modes, and works with TMT and dimethylation labeling of N-termini. Similarly, users that work with different labeling strategies (e.g. acetylation or iTRAQ) can either replace the modifications with strings matching CLIPPER 2.0 standards, or directly modify the underlying patterns in the code to match their modifications (in clipper.py, get_patterns_* functions).

### Processing steps and database annotation

We make use of the argparse module of the standard python library^19^ to pass user specified arguments to the tool. After a format check for the input table, data columns and rows, we infer the software used if it is not specified by the user. Initially, the input table is checked for rows with empty accession numbers, as well as empty sequences and invalid protein alphabet characters. By default, complete annotation is performed on all peptides, but sequences that are not N-termini can optionally be removed for faster result turnover. Remaining peptides are then annotated using the Expasy and SwissProt modules in the Biopython^20^ package to dynamically query the Uniprot database API^21^ for an entry matching the corresponding protein accession. The gene name, full sequence, protein description, as well as gene ontology information and known processing events are retrieved and matched to the peptide. The peptide sequence is mapped to the protein sequence to annotate peptide starting and ending positions, the p1 cleavage site position, and the cleavage site environment with a user specified length. The concurrent.futures^19^ module is used to offer threading capabilities when retrieving UniProt entries, enabling substantially faster annotation in multi-core computer architectures. Users can adjust the rate of Uniprot requests with an argument.

Static snapshots of ProteinAtlas^22^, MEROPS^23^ and AlphaFold^24^ databases are used in CLIPPER 2.0, to optionally perform further annotation. Snapshots of ProteinAtlas and MEROPS are included, but due to the large size of the Alphafold database, it should be separately downloaded by the user, and the path should be specified by setting “alphafold_folder_name = “ equal to the local Alphafold path in annutils.py. ProteinAtlas is used to map UniProt entries in the dataset to chromosome location, RNA expression in cell types and tissues, protein class, disease involvement, and gene ontology information for each protein. We use the Alphafold database to load model structures for each protein in Pymol^25^, and calculate secondary structure prediction for the cleavage sites in the dataset using the ‘dss’ function from the Pymol command line, and the solvent accessible surface area with the ‘get_area’ function. Finally, we report known proteases either in Uniprot or the MEROPS database for the cleavage site. The annotation columns are inserted directly in the input table by default, with the option to save the annotation file separately if preferred. A description of additional columns generated with CLIPPER 2.0 is available in the tool repository.

### Proteoform certainty and protease activity prediction

We calculate proteoform certainty based on the number of protein accessions containing the sequence of the identified peptide, with certainty as the reciprocal of accessions (1/N, where N= number of proteins containing that peptide). For aminopeptidase activity prediction, we sort all identified peptides by length, and match peptides that share the last five residues in their sequence. For the matched peptides, we check whether the parent peptide differs only by one or two residues in the N-terminal side, in which case they are annotated as aminopeptidase or dipeptidase activity, respectively. The annotated peptides become parent peptides in the next round, and the process is continued until there are no peptides with ragging patterns compared to their parent peptides.

We also provide a generic module for protease specificity matching and substrate suitability prediction, using MEROPS data and position specific scoring matrices (PSSMs). CLIPPER 2.0 accepts a user input file with MEROPS protease identifiers (one identifier per line). For each protease of interest, as supplied in a .txt file with one MEROPS identifier per line, we use its substrates in MEROPS to construct a PSSM^26^ matrix encoding the protease sequence specificity motif. This is then used to score all identified cleavage sites and provide a summed log odds ratio score across all positions from the annotated cleavage site. The user has the option to use the Blosum62 peptide alphabet^27^ to add pseudocounts to the PSSM, which can be particularly useful if there is a limited amount of substrates in the MEROPS database.

### Statistical analysis

To perform statistical analysis with CLIPPER 2.0, the user has to provide a condition file in addition to the peptide report. The file consists of one condition per line, initiated by a condition name which is followed by space separated strings that are unique to a quantification column associated with the given condition. These could be TMT reagents names if present, but any unique substring can be used. Before performing statistical analysis, we match the quantification columns specified by the user in the condition file, and convert the data to numeric type. We also provide the option to fill missing values with a numeric value of choice or remove them. While supplying a condition file is not mandatory for CLIPPER 2.0 to run, it is necessary for statistical analysis and therefore for most figure generation.

To designate peptides in the ends of the fold change distribution, we calculate the density function from the histogram of fold change values using numpy, and classify peptides as “significant high” or “significant low” in the distribution.

We perform statistical testing across conditions using the statsmodels package^28^ and the ttest_ind, or scipy^29^ and the f_oneway function for student’s T-test or analysis of variance (ANOVA), respectively. The user has the choice to perform pairwise t-testing instead of ANOVA (which is the default option if more than two conditions are present). We also use the multitest module to perform multiple testing correction, which is recommended when performing statistics in large datasets. Per default, corrected p-values are reported in the output file, but are not used for figure generation. This can be changed by supplying the -mt argument at runtime, and the type of multiple testing correction can be specified with -mt argument.

### Visualization and enrichment analysis

CLIPPER 2.0 uses the matplotlib^30^ and seaborn^31^ libraries for visualization. Seaborn is used to create a barplot of identified and quantified peptides, and scatter plots with user defined cutoffs to create volcano plots of log2(FC) and -log10(p-value) for each comparison of conditions in a pairwise manner. The CV distribution is visualized by a kernel density estimation with kdeplot, and histplot generates a histogram showing fold change distributions between conditions. Heatmap and clustermap functions use euclidean clustering and z-score distribution to visualize the dataset and provide a way to assess consistency across conditions. Pie charts are generated with matplotlib pie function to visualize the relative number of different types of N-termini. Dimensionality reduction algorithms are used with sklearn^32^ to perform principal component analysis (PCA) and visualize the first two principal components and their encoded variance, as well as a Uniform Manifold Approximation and Projection (UMAP)^33^ plot. We use matplotlib and Pymol to visualize protein cleavages on AlphaFold protein structures with a per protein normalized color scale indicating fold changes for each peptide, as well as DNA Features Viewer^34^ for sequence visualization and domain highlighting. We utilize the gProfiler^35^ python interface to perform overrepresentation tests, and plot significant terms per condition, with a capped number of terms (30) per plot. Finally, we use the Reactome^36^ API and the reactome2py wrappers to carry out pathway analysis, and NetworkX^37^ for graph arrangement and visualization.

Logo plots are generated based on the cleavage sites with a default but adjustable size of p4-p4’. The provided sequences are used to generate count and frequency matrices, and the frequency matrix is optionally corrected with Blosum62 pseudocounts. A weight matrix is found as in eq. 1, where *w*_*a*_is the weighted value for each amino acid at a position, *p*_*a*_ is the frequency of the amino acid, and *q*_*a*_ is its background frequency. Probability logos are created based on the frequency matrix and PSSM logos based on the weighted matrix. Shannon information logos^38^ are created by calculating the information content of each amino acid at each position based on the frequency matrix, assuming equal amino acid background distribution, while the Kulback-Leibler divergence^39^ corrects the information content based on the background amino acid distribution of sequences in TrEMBL release 2023_05.

## Results & Discussion

### Annotation of cleavage environment and protein information

CLIPPER 2.0 takes in the peptide report from supported proteomics database search engines, without a need for the protein table (Suppl. Fig 1). To annotate the peptides sequences, our algorithm matches the protein accession to a uniprot entry, searches the protein sequence for the peptide, and annotates the peptide location, as well as the cleavage environment.

Beyond the cleavage annotation, we also retrieve general information for the protein, in particular gene name, description, and protein length. Notably, CLIPPER 2.0 utilizes the information available in the Uniprot entry to match and annotate the cleavage with known processing events, such as signal, transit, or propeptide removal, allowing classification of unknown sites as neo N-termini.

### Prediction of protease cleavage events

In addition to general information and cleavage environment, CLIPPER 2.0 annotates known protease cleavages based on entries in Uniprot and MEROPS, providing protease names and MEROPS identifiers for proteases with known substrate cleavages identified in the dataset. We also employ information available in Protein Atlas to retrieve and report on protein expression in different tissues and cell types (only available for human and mouse proteins), chromosome location, protein class, and gene ontology associated with the protein entry. Furthermore, we perform a basic calculation on the number of proteins associated with the given peptide sequence to report on the proteoform certainty of the identified cleavage, and ragging peptides are identified as aminopeptidase activity.

If a specific protease is under study, CLIPPER 2.0 allows the inclusion of a .txt file containing line separated MEROPS protease identifier(s). Based on prior substrate sequence knowledge in MEROPS, CLIPPER 2.0 generates a position specific scoring matrix (PSSM) and calculates a likelihood score for each cleavage site, which indicates the cleavage similarity to known motifs.

CLIPPER 2.0 uses AlphaFold models along with the PyMOL engine to provide a secondary structure prediction for the immediate cleavage environment, and designates each amino acid surrounding the cleavage site as either part of a loop (L), helix (H), or beta strand (S). A value for solvent accessibility is also computed in square Angstroms and provides a measure of cleavage site exposure.

### Statistical analysis of cleavages

CLIPPER 2.0 optionally accepts a condition file which specifies the conditions used in the experiment, and information about individual replicates. These quantification columns are used by the tool to perform statistical analysis of peptide abundances (which is intended but not restricted to terminal peptides), and report several quality control and significance metrics. We compute mean, standard deviation and coefficient of variation (CV) for conditions included in the experiments to allow for quick filtering and quality control of the experiment along with general dataset statistics (Fig. 2a-b). If more than two conditions are present for an experiment, users can choose between ANOVA or pairwise two sample T-test, otherwise a T-test is used for statistical analysis. CLIPPER 2.0 then reports fold change, log2 fold change, p-value, and multiple testing corrected p-value for each peptide tested. Per default, multiple testing corrected values are included in the output table, but are not used for figure generation. For cases where statistical testing is not optimal, we also compute the distribution of fold changes across conditions and annotate peptides in the high and low 5% tails of the distribution as “significant”.

**Figure 2.**
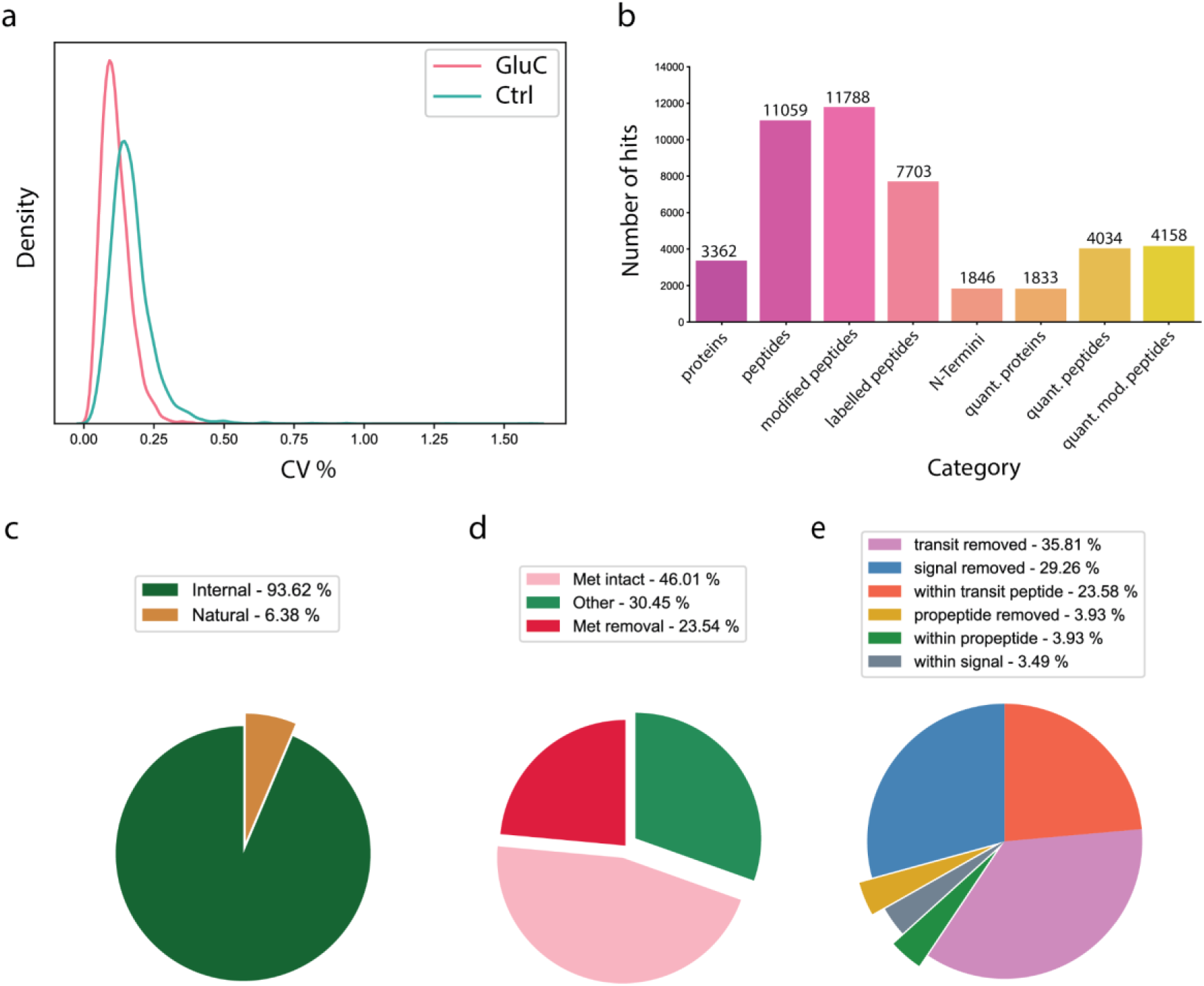
GluC digested HeLa lysate followed by TAILS. General data QC plots are shown. a) Coefficient of variation distribution as kernel density estimation plot. b) Bar plot of identified and quantified proteins, peptides, and N-termini. The number of unique peptide sequences, quantified proteins and peptides are also reported. c) Pie charts illustrating number of natural (Met-intact, Met-removed, and other known processing events) or internal N-termini. d) Pie chart showing percentages of natural N-termini with Met-intact, Met-removed or other natural cleavages. e) Pie chart showing annotated known processing events found in relevant databases.

### Visualization of dataset and analysis

A main component of CLIPPER 2.0 is its visualization capabilities. By default, pie charts are generated for the different categories of N-termini (natural, signal peptide removal, and others, Fig. 2c-e). Additionally, a bar plot showing the number of proteins, peptides and N–termini, along with modified and labeled peptide numbers, is generated for a check of experiment quality.

If the user has provided an input table with quantification values and a descriptor file detailing the conditions used in the experiment, several plots are generated for quality check and dataset exploration. Toward that end, the coefficients of variation across replicates for each condition are also displayed as a density distribution plot. To explore data quality and replicate similarity, a heatmap is also generated. To observe similarities across conditions and replicates, CLIPPER 2.0 generates a clustermap plot, along with PCA and UMAP plots (Fig. 3a-b).

**Figure 3.**
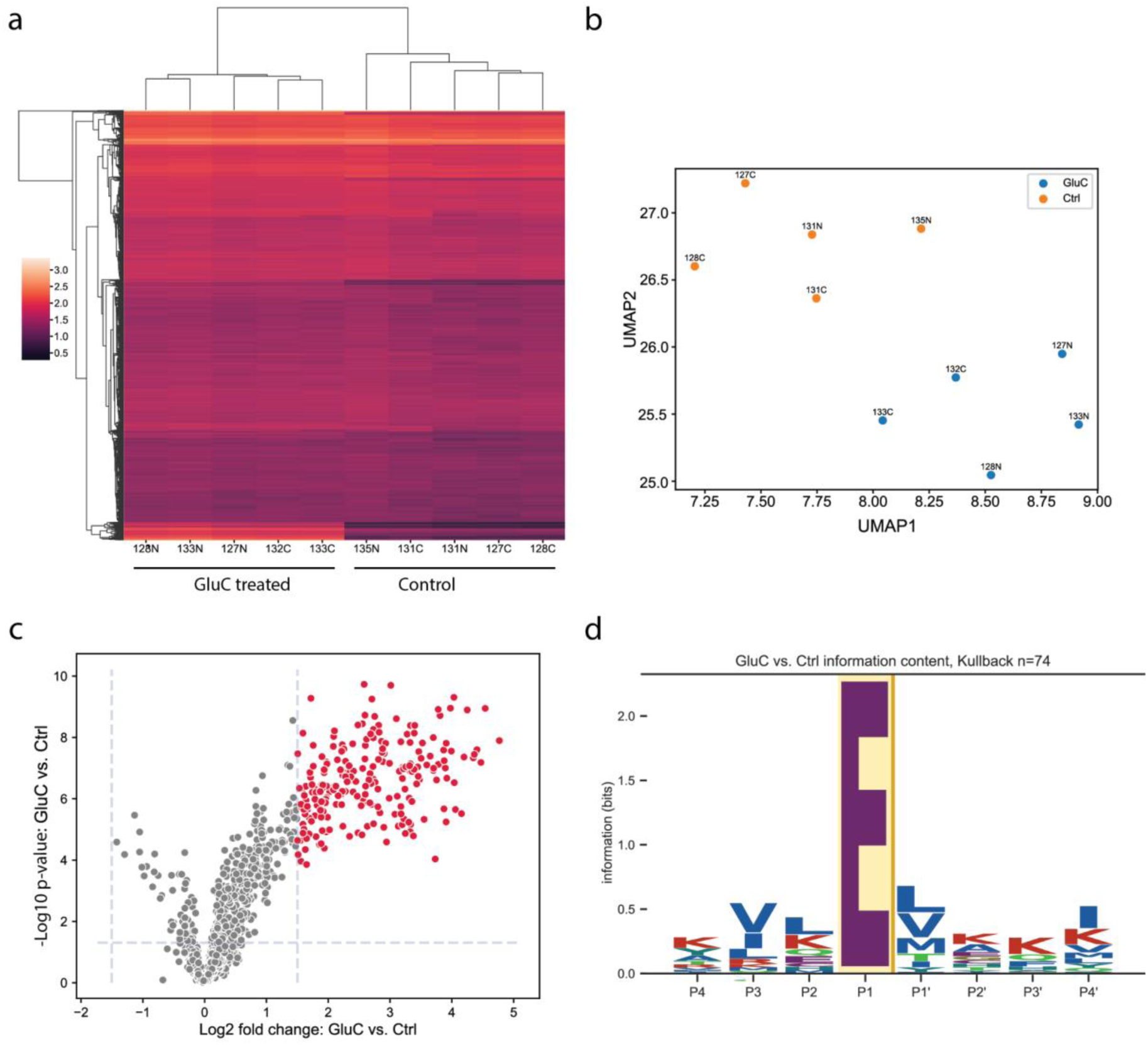
a) Clustermap including replicates of all experimental conditions and their similarity across replicates (columns) and identified peptides (rows). b) UMAP representation of replicates across all conditions. c) Volcano plot showing log2 transformed fold change values for peptides in the dataset, and the -log10 p-value from the statistical testing performed. Significant peptides according to cutoffs provided by the user are highlighted in red. d) Sequence motif specificity for the significant cleavages observed in a pairwise comparison of conditions used in the experiment.

Beyond the general dataset descriptive plot, CLIPPER 2.0 visualizes the statistical analysis performed with fold change distribution and volcano plots (Fig. 3c). Fold change ratios for all pairwise comparisons of conditions in the experiment are visualized with a histogram and kernel density estimation in the same plot (Suppl. Fig. 2a), allowing for selection of relevant significance cutoffs. For the volcano plots, cutoffs are based on empirically observed values and can be modified by the user, with significant peptides displayed with different hue.

Sequence motif logos are widely used to visualize protein specificity in e.g. transcription factor binding motifs and kinase specificities. Similarly sequence logos allow for facile and intuitive visualization of protease cleavage specificity, in a more simplified way compared to specificity heatmaps showing all amino acids across positions. For that reason, we opted for visualizing cleavage specificity with sequence logos (Fig. 3d), using the cleavage environment annotation in previous steps. The logos can be computed with several established methods, such as PSSMs (Suppl. Fig. 2b) or Kullback–Leibler divergence matrices. For each condition tested, peptides which are either significantly higher or lower in abundance and their cleavage sites are used to construct the logos specified by the user, with the number of sequences and the method used for logo generation displayed above the plot.

In order to directly visualize differences in significant N-termini across conditions, a gallery with bar plots showing abundances for significant N-termini and the natural N-terminus of the protein of origin is also generated (Suppl. Fig. 3). This allows for direct comparison of natural and internal cleavage abundances, and their changes across conditions at the protein and peptide level.

Significant cleavages can also be visualized in the protein sequence, annotated with protein– wide normalized fold changes across conditions. If relevant post translational processing information is available for a protein in the Uniprot database, these regions will also be included and labeled in the sequence plot. Similarly, cleavages by known proteases either in MEROPS or Uniprot databases will also be included in the plot . This allows for quick overview and exploration of experimental cleavages, and facile and extensive comparison with known cleavages. In addition to sequence visualization, CLIPPER 2.0 uses the AlphaFold model database to retrieve, label and visualize experimental cleavages and fold changes across conditions directly on the protein structure (Fig 4a). This feature enables examination of observed cleavages on different regions of proteins, and can be useful when examining cleavages that are solvent accessible or in different secondary structures.

**Figure 4.**
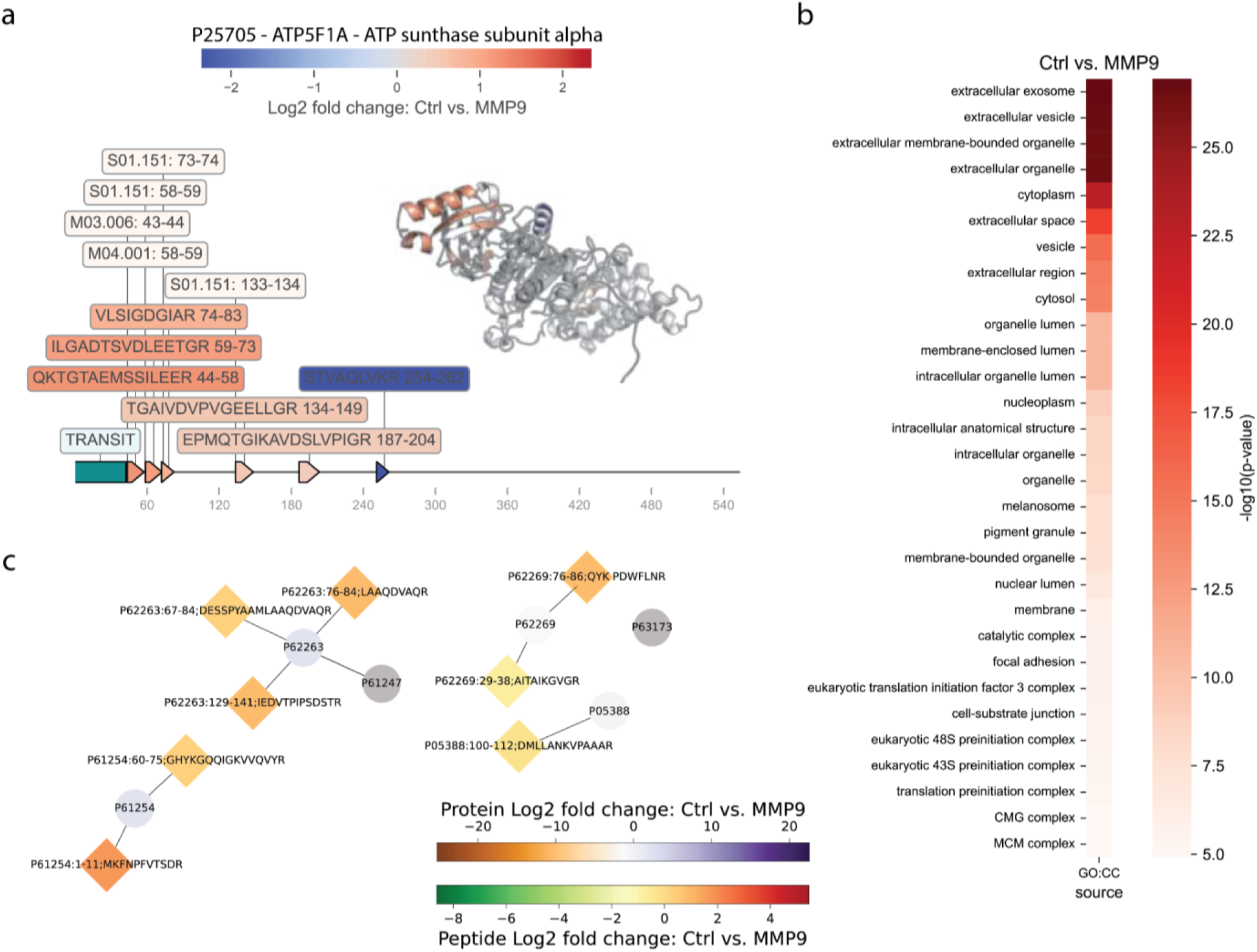
a) Sequence and structure visualization of peptides and annotation of known protease cleavage sites in ATP synthase subunit alpha. b) Functional enrichment reveals a preference for extracellular proteins, corresponding with the role of MMP9 as an extracellular protease. c) Pathway visualization for R-HSA-1799339: SRP-dependent cotranslational protein targeting to membrane. Selected parts of the pathway are shown with known interactions between proteins, and the cleavages observed. Identified cleavages are represented as diamonds, whereas proteins are shown as circles. Fold change scales are shown separately for protein and peptide levels.

### Network analysis and functional enrichment

We use gProfiler to perform gene set enrichment analysis for the significantly differentially abundant proteins in each condition. As with the majority of illustrations, this is only possible if the user has assigned conditions and performed statistical analysis. CLIPPER 2.0 extracts significant terms for each condition, and exports visualizations optimized for clarity and robustness. With this, users obtain functional enrichment results for their experimental datasets, which can be explored and interpreted intuitively (Fig. 4b). In addition to the functional analysis of our tool, we use the Reactome web services and available python wrappers to perform network analysis for differentially abundant proteins, and extract significantly enriched pathways for subsequent visualization (Fig 4c). We calculate and show fold changes between conditions in the resulting plot, which are also annotated with the observed cleavages and their positions for the proteins of the dataset. Both enrichment and pathway plots only work with human datasets per default.

### Future implementations

CLIPPER 2.0 will be regularly maintained and input for improvements will be welcome. Future versions are planned which will incorporate a general input format for search engine outputs which currently are non-supported, support for C-terminomics analysis, and compatibility with improved pathway analysis tools by overlaying data on known annotated pathways from e.g. KEGG and WikiPathways. Protease prediction in CLIPPER 2.0 is still rudimentary due to the lack of protease cleavage specificity information, and can be improved by utilizing generated datasets instead of relying solely on database information. The data analysis done for degradomics has many parallels to that for other PTMs, as much work is done on the peptide level, importance is placed on specific positions within the sequence, and comparisons are often performed between modified and unmodified forms. As such, CLIPPER 2.0 can be expanded to fit other purposes than degradomics analysis.

## Conclusion

Here, we introduce CLIPPER 2.0, a robust, scalable, light-weight and user-friendly tool for comprehensive data analysis of degradomics experiments utilizing different workflows and proteomics search engines. Our tool makes significant strides towards automation and complete end-to-end pipelines for degradomics experiments. We describe several improvements over state-of-the-art software packages by employing statistical analysis, advanced annotation with available databases and visualizations to help with initial data exploration.

In summary, CLIPPER 2.0 improves both annotation, computation, parallelism, and publication ready visualizations for fast analysis of large datasets. We streamline annotation and analysis of degradomics dataset in a manner compatible with automated workflows and high throughput analysis. Finally, we will regularly update CLIPPER 2.0 based on community feedback, and hope for quick adoption of the tool in various experimental workflows.

## Data Availability

CLIPPER 2.0 can be accessed on GitHub (https://github.com/UadKLab/CLIPPER-2.0). The datasets used in this study have been deposited to ProteomeXchange Consortium^40^ via the PRIDE database^41^ and are available with the dataset identifier PXD047261. The reviewer account has the following credentials: username -, password - CdYg44tR.

## Contributions

KK and UadK conceived the research idea. KK and AMH designed and programmed the tool with input from RH, EM, EMS and UadK. KK, AMH and EM selected the validation datasets. KK, ADL and EM performed sample preparation and mass spectrometry analysis, and KK, AMH and EM analyzed the datasets. KK and AMH wrote and edited the manuscript with input from all authors. All authors read and approved the final version of the manuscript.

## Acknowledgements

EMS and UadK acknowledge support by a Young Investigator Award from the Novo Nordisk Foundation (NNF16OC0020670) and PRO-MS: Danish National Mass Spectrometry Platform for Functional Proteomics (grant no. 5072-00007B).

## Supplementary figures

**Supplementary figure 1.**
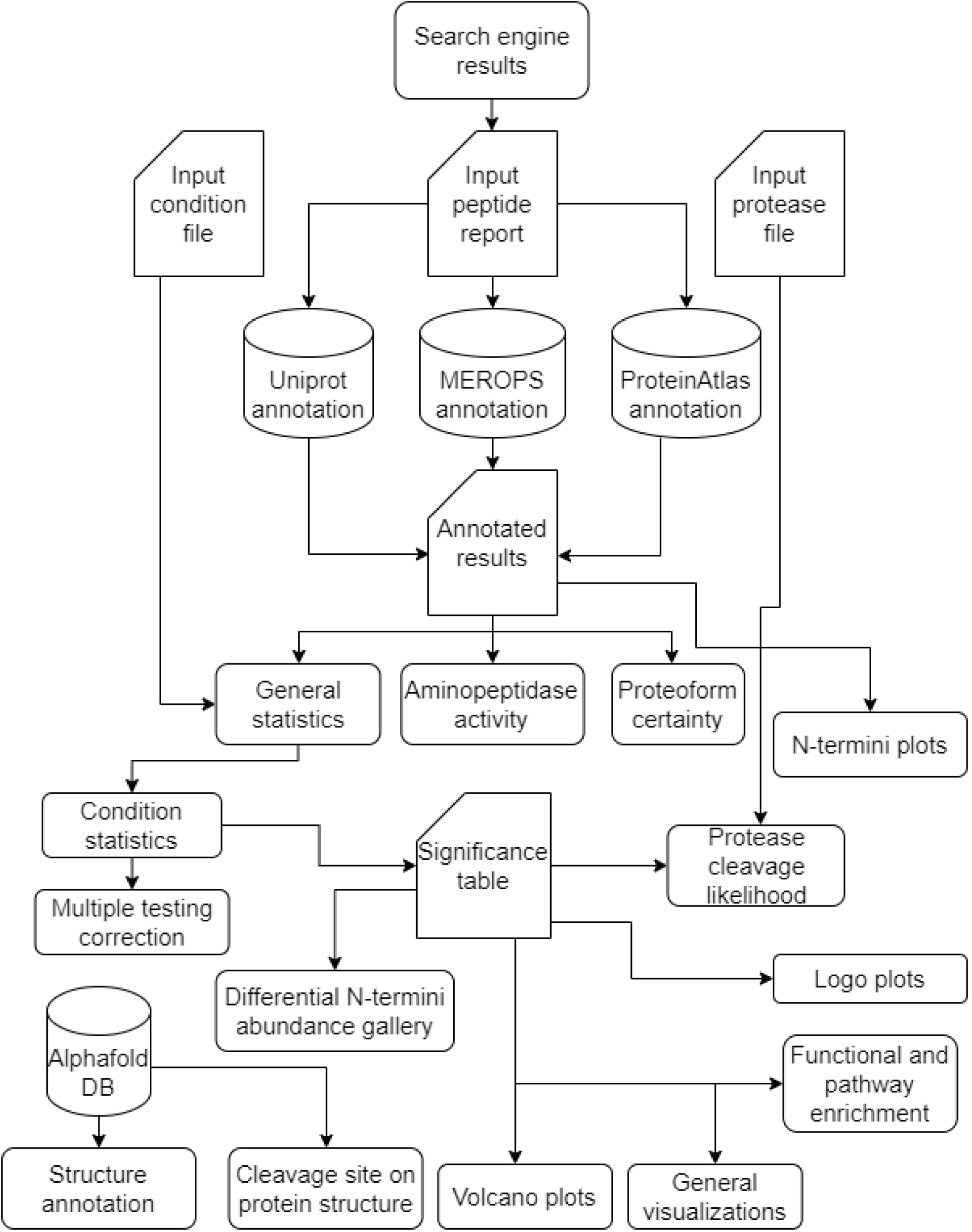
CLIPPER 2.0 flow chart indicating different features, capabilities and results generated.

**Supplementary figure 2.**
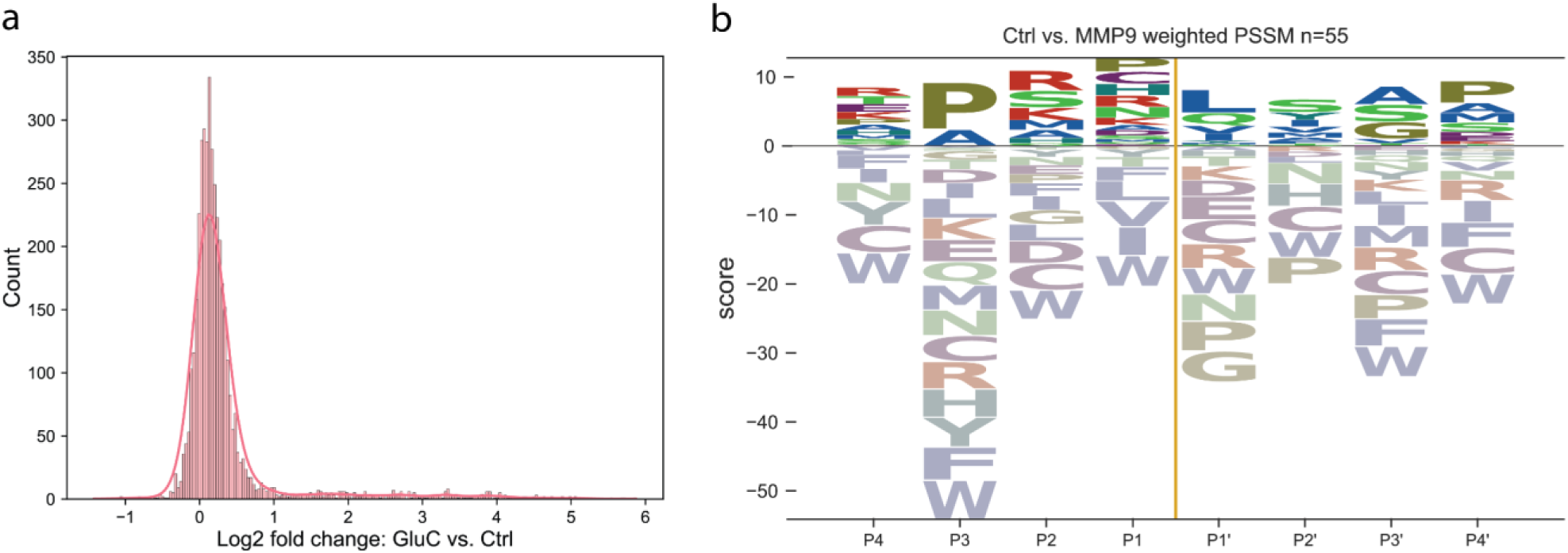
a) Fold change histogram distribution with KDE estimation. b) PSSM values for MMP9 specificity visualized as a cleavage logo.

**Supplementary figure 3.**
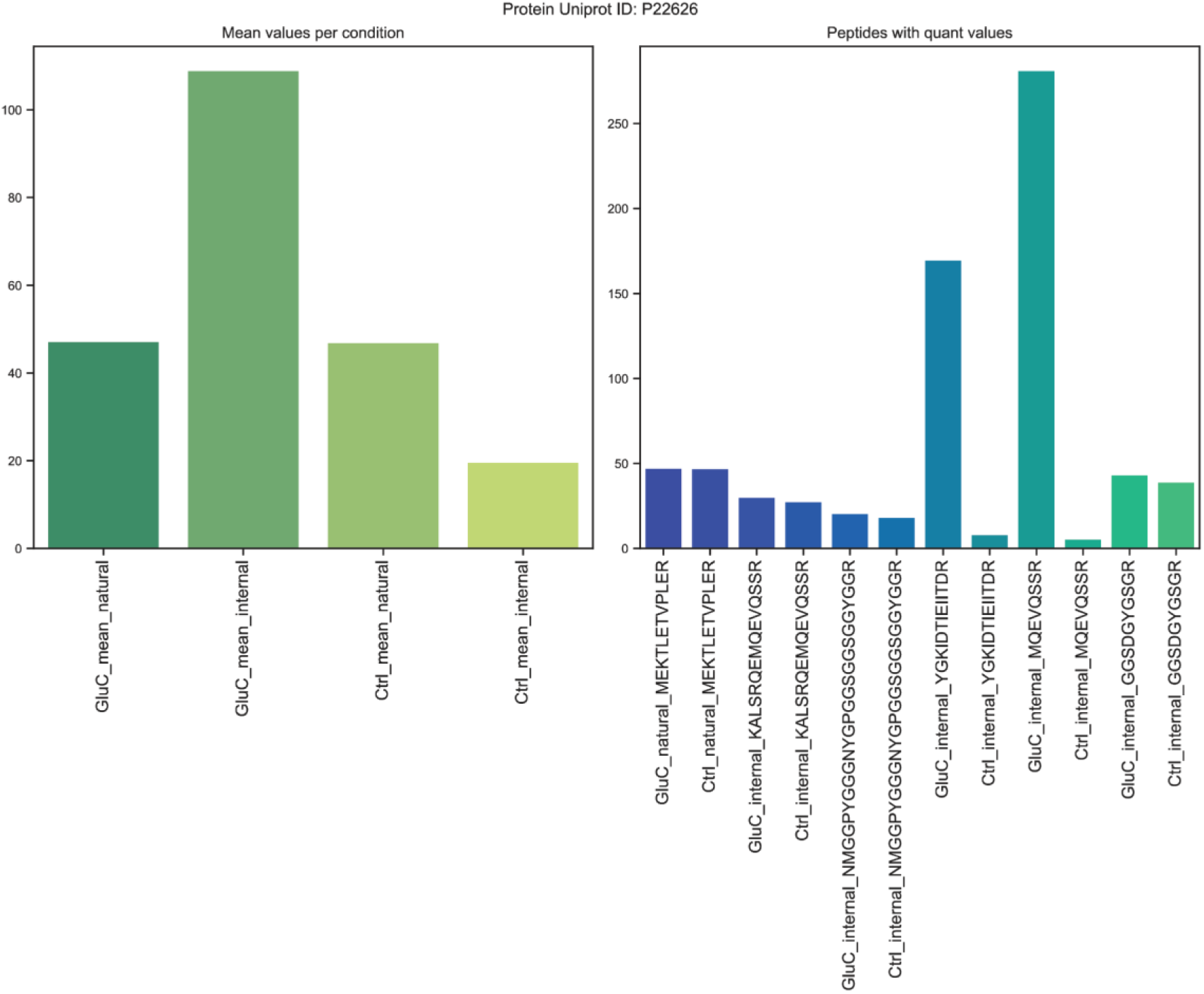
Natural and internal peptide abundance for termini identified in HNRNPA2B1 (Uniprot ID: P22626) as a mean abundance (left), and for each individual peptide (right).

